# SURFR: genome-free discovery of cancer-unique small RNAs from cohort sequencing data

**DOI:** 10.1101/2025.10.07.680618

**Authors:** Panagiotis Kalogeropoulos, Konstantin Danilov, Maria Alejandra Ulloa, Asimina Talanti, Bastian Fromm, Simon Ekman, Per Hydbring, Marc R. Friedländer

## Abstract

Small RNAs play numerous roles in cancer biology and function as biomarkers and therapeutic targets. A recently discovered class of small RNAs - the orphan non-coding RNAs (oncRNAs) - are uniquely present in cancer and absent from healthy tissues. Here, we developed a reference-free computational approach that is independent from the human genome and applied it on cohort sequencing data from five cancer types to discover 1211 novel oncRNAs. Most of these new sequences do not map perfectly to the human genome, and hundreds correlate with patient outcomes. Our new oncRNAs include the first bona fide cancer-unique microRNA; oncRNAs predictive of patient survival and a small RNA that is expressed from an evolutionarily ancestral part of the genome that has been lost in most human individuals. In summary, we provide our reference-free computational approach and more than thousand new cancer-unique RNAs as resources to the community.

## Main

Small RNAs are non-coding RNA molecules shorter than 200 nucleotides that can dynamically regulate gene expression and perform house-keeping functions and have been implicated in numerous biological processes including cancer^1–3^. Among these, microRNAs comprise the most extensively studied class and are known to function as both oncogenes and tumor suppressors^4,5^. In addition, other classes of small RNAs, including small nucleolar RNAs (snoRNAs), transfer RNAs (tRNAs), small nuclear RNAs (snRNAs), and PIWI-interacting RNAs (piRNAs), have gathered attention for their roles in tumorigenesis^2^.

More recently, Goodarzi and colleagues described a novel class of small RNAs, termed orphan non-coding RNAs (oncRNAs), which are specifically expressed in cancer and absent in healthy tissues. They provided evidence that one such cancer-unique oncRNA, derived from the TERC transcript, contributes functionally to breast cancer metastasis^6,7^. Moreover, they have shown that these molecules may serve as potential cancer biomarkers from liquid biopsies^8^.

Next-generation sequencing (NGS) enables highly sensitive detection of small RNAs^9^, however, the novelty of oncRNAs and the fact that most of them likely remain undiscovered means they are not included in existing annotations. This makes them in effect invisible to conventional computational approaches that rely on known transcript databases. Furthermore, the genomes of cancer cells can highly deviate from their healthy counterparts due to large structural variations and small sequence variants caused by the disease^10–13^. At the same time, the genomes of individual humans also diverge from the human reference genome^14^. These complexities underscore the need for unbiased, reference-free analytical methods to identify novel small RNAs such as oncRNAs in cohort sequencing data.

In this study, we substantially expand the catalogue of oncRNAs by discovering more than thousand new cancer-unique molecules and characterizing their landscape across five distinct cancer types. We present a novel computational pipeline that bypasses the need for both genome alignment and pre-existing annotations. Departing from traditional workflows that map reads to the reference genome and quantify only annotated transcripts, our method directly quantifies and compares raw small RNA sequences, drawing inspiration from recent reference-free computational advances in both mRNA and small RNA analysis provided by the SPLASH2^15,16^ and Seqpac^17^ algorithms respectively. Applying this approach to publicly available sequencing data from the TCGA^18^ and CPTAC^19^ cohorts, we discover 1211 previously unannotated oncRNAs and highlight a subset of these molecules of particular interest.

## Results

### The genome reference-free SURFR algorithm

The most common methods to discover cancer-associated transcripts from sequencing data are feature-based^17^. In these approaches, the sequenced RNAs are mapped to a reference genome and overlapped with gene annotations (features). This approach works well when the aim is to quantify previously annotated molecules. However, we anticipated that orphan non-coding RNAs (oncRNAs) may arise from the transcription of normally silenced regions in cancer and therefore do not overlap with any known or annotated genomic features (Fig. 1a). In addition, transcripts that are unique to cancer may originate from genomic aberrations that do not resemble sequences in the reference human genome^10–12^, and therefore cannot be mapped. Therefore, the traditional feature-based approaches might not be suitable for detection of oncRNAs. We addressed these issues by using a reference-free and sequence-centric approach that treats each unique sequence as an independent entity, regardless of its genomic mapping or annotation status. Hence, our method generates tables that sum up counts for distinct sequences rather than features (Fig. 1b).

**Figure 1:**
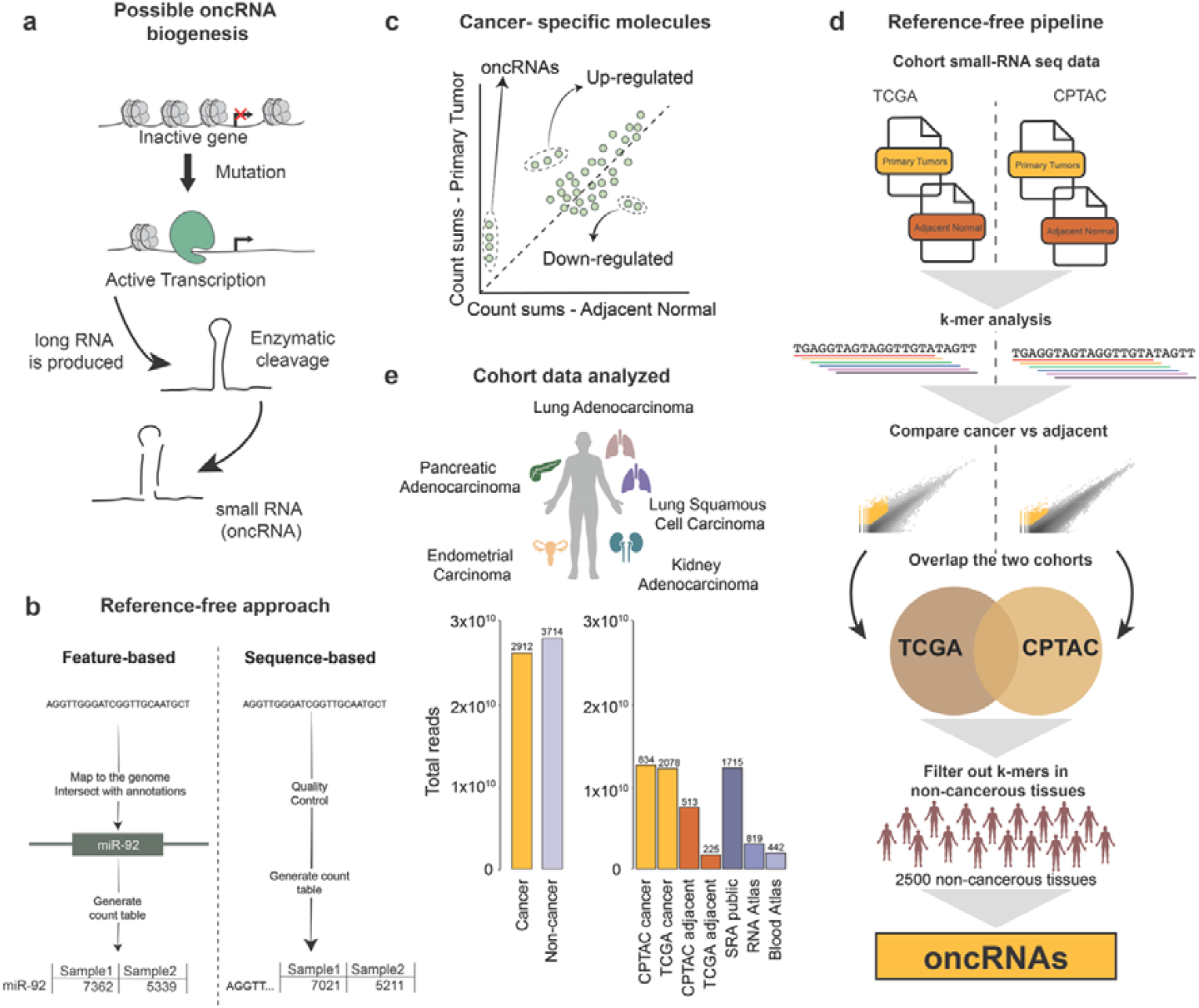
oncRNAs and the reference-free SURFR algorithm. **a**, Example of possible oncRNA biogenesis by a mutation leading to activation of transcription and further processing by the small RNA machinery. **b**, oncRNAs are identified as the sequences that are expressed in cancer samples but have near-zero expression in non-cancerous tissues. **c**, The SURFR pipeline, small RNA sequencing data from two cohorts are split into k-mers of length 17. The k-mer counts are aggregated across samples of the same condition and compared, k-mers that show high cancer specificity are selected for further analysis. The selected k-mers from each of the two cohorts are intersected and their expression checked against a cohort of >2,500 non-cancerous human tissue samples to verify cancer-specificity. Finally, the k-mers are concatenated into contigs which are the final set of oncRNAs. **d**, SURFR uses a reference-free approach, meaning that reads are not mapped to the reference genome but rather each sequence is considered a separate entity for quantification. **e**, The data that were analyzed in the study and the distribution of reads and samples across cohorts.

This computational methodology for Sequence-based Unbiased Reference-Free small-RNA sequencing analysis that we call SURFR, consists of the following core steps: (a) in each condition the raw Fastq files are pooled together; (b) the pooled files are split into k-mers and (c) the k-mer counts are compared between the conditions (Methods). This unbiased approach can be used to compare the expression of small RNAs in any setup where two or more conditions exist. In the context of oncRNAs, we compared the expression counts of 17-mers between tumor samples and adjacent non-cancerous tissues. The oncRNAs, by definition of their cancer-uniqueness, are the sequences that are detectable in cancer but practically absent in non-cancer samples (Fig. 1c).

Our input data consisted of small RNA sequencing data from The Cancer Genome Atlas (TCGA)^18^ and the Clinical Proteomic Tumor Analysis Consortium (CPTAC)^19^ consortia, focusing on primary tumors and adjacent normal tissues (Methods) across five shared cancer types: Lung Adenocarcinoma (LUAD), Lung Squamous Cell Carcinoma (LUSC), Kidney Adenocarcinoma (KIAD), Pancreatic Adenocarcinoma (PAAD) and Uterine Corpus Endometrial Carcinoma (UCEC) (Methods). This encompassed over 3,500 samples and over 30 billion reads, with a skew towards cancer samples (Fig. 1d). Following quality control, we aggregated reads within each condition and segmented them into 17-mers (Methods). We then compared the abundance of each 17-mer between tumor and adjacent samples, retaining only those significantly enriched in tumors for further analysis. This process was performed independently for TCGA and CPTAC, and only 17-mers that were significantly enriched in both cohorts were retained. Finally, to stringently ensure true cancer uniqueness, we quantified the remaining 17-mers across a diverse panel of over 2,500 public human non-cancerous tissues samples, including various healthy tissues and non-cancerous disease states (Fig. 1d, Fig. 1e, Methods, Supplementary Table 1). Sub-sequences that were present in the non-cancer control samples were deemed to fail to demonstrate cancer uniqueness and were excluded (Fig. 1e). The inclusion of samples from infections and immune cell types in these control samples served as a robust measure to distinguish cancer-unique 17-mers from those potentially derived from inflammation. Lastly, the cancer-unique 17-mers were concatenated into larger sequences and mapped against known artificial sequences to remove any remaining sequencing artifacts (Methods). No matches to adapters or primers were found, suggesting that the two-cohort approach successfully removed the artifacts.

### SURFR discovers 1211 novel oncRNAs in five cancer types

We applied SURFR to the five cancer types mentioned above and identified 1211 oncRNAs (Supplementary Table 2). The number of oncRNAs discovered ranged greatly, from 12 in pancreatic adenocarcinoma to 816 in endometrial carcinoma (Fig. 2a). In general, the results highlight the importance of using a two-cohort approach to narrow down the number of candidate molecules and remove sequencing related artifacts and false positives. This appeared to be especially important when there is an imbalance between the number of cancer and control samples in a cohort. For instance, the TCGA lung adenocarcinoma cohort has more than 10 times the number of tumor samples compared to adjacent samples, whereas CPTAC has almost an almost equal number of tumor and adjacent samples for this cancer type (Fig. 2b). This imbalance results in a 100-fold more 17-mers that passed the first filtering step in TCGA (n = 240,843) compared to CPTAC (n = 2,466) (Fig.2c). Out of those however, only 293 were actually present in both cohorts (Fig. 2d). While this stringent filtering step can lead to loss of some genuine cancer-specific molecules, it greatly improves specificity (see Discussion section). Next, sub-sequences with high expression in the public non-cancerous cohort were also filtered out (Fig. 2e). The cancer expression pattern of these 17-mers was highly variable, with cancer counts summed over all cancer samples ranging from 200 counts up to 10,000 while maintaining the strict criteria of low counts in non-cancerous tissue (Fig. 2f). As a final step, 17-mers with overlapping sequences were combined into larger contigs, resulting in a pool of 73 distinct sequences, each of which is an oncRNA.

**Figure 2:**
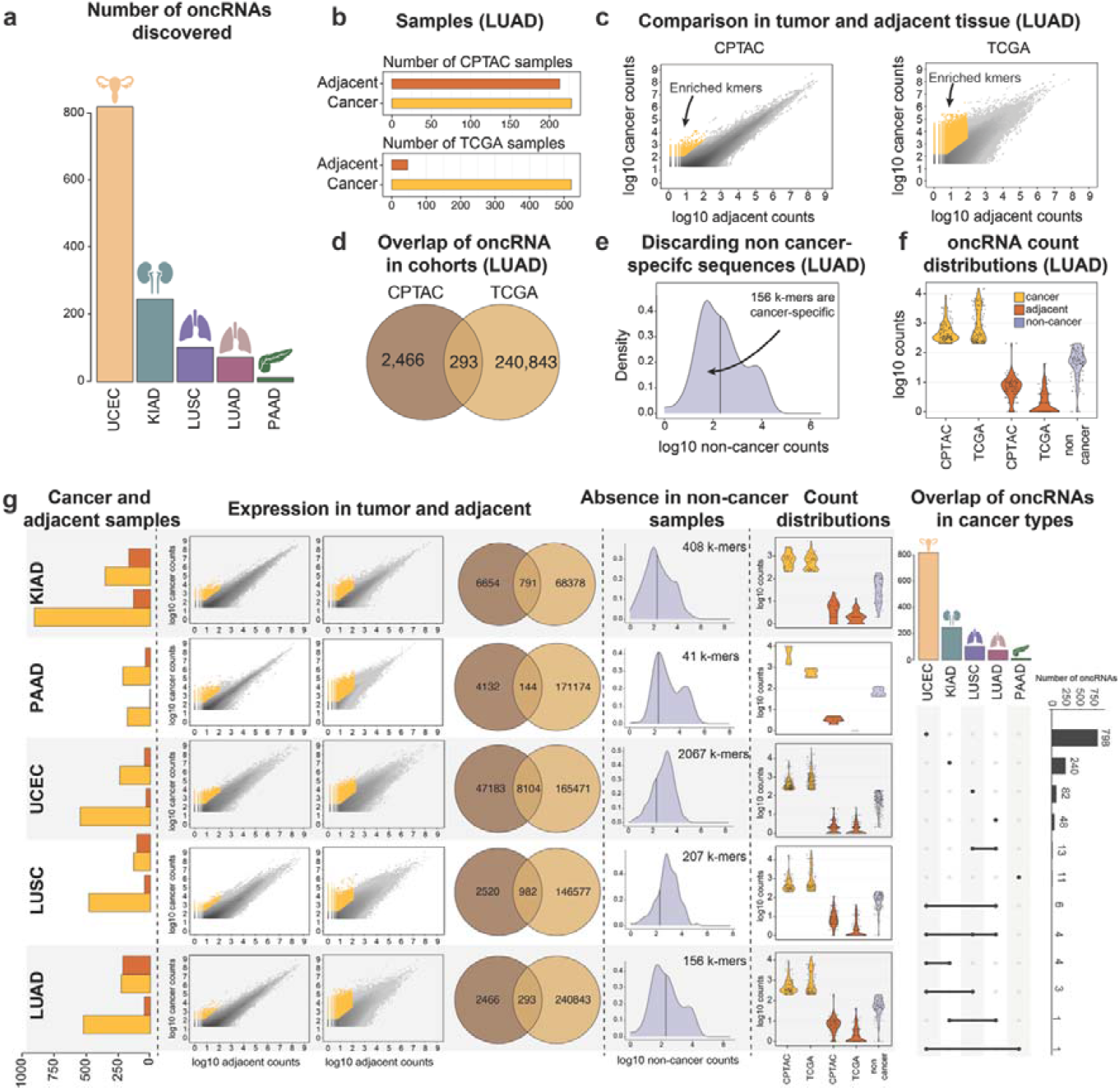
The oncRNA landscape across five cancer types. **a**, The total number of oncRNAs discovered by cancer type. **b**, Numbers of samples of lung adenocarcinoma (LUAD) across the two cohorts. **c**, Distribution of all k-mer counts and selection of the cancer-specific ones. **d**, Cancer-specific k-mers in each cohort and their overlap. **e**, 17-mers that have more than 200 sequence counts in the non-cancerous cohort are filtered out. **f**, Counts of the cancer-unique k-mers that have passed all the filtering criteria across cohorts. **g**, Similar to B-F but for all cancer types and the overlap of oncRNAs between cancers.

The same approach as described above for lung adenocarcinoma was performed for the other cancer types (Fig. 2g). We then investigated how many of these sequences overlap between cancer types. In general, we do not see high overlap between cancer types, suggesting that most of the oncRNAs are also specific to a given cancer. Naturally, the highest overlap is found between the two non-small cell lung cancers, with 13 sequences in common. Overall, we are able to identify many oncRNAs that appear to be mostly specific to a cancer type (Fig. 2g). An important advantage of using SURFR for the analysis is that we are not limited to sequences that are present in the human reference genome and can capture canonical small RNAs harboring point mutations, microRNA variants^20^ and also sequences completely missing from the reference genome. To highlight this, we mapped the oncRNAs to the human genome and found that only 512 (42.1%) of all oncRNAs mapped without nucleotide mismatches, while 620 (51.2%) mapped with a single mismatch (Extended Data Fig 1b). Of the oncRNAs that could be traced to the human genome, 450 originated from long-noncoding RNAs (lncRNAs). These cases may be similar to mature microRNAs that are cleaved out from longer non-coding precursor transcripts by endonuclease action^21^. Another 409 oncRNAs originated from exons of protein coding genes. It has previously been described that small RNAs can be cleaved out from mRNAs^22^, leading to the production of a short transcript and the destruction of the longer one. Interestingly, 81 (6.7%) of the oncRNAs did not map to the human reference genome at all, suggesting that they could be the result of cancer-related genome aberrations. These percentages were overall similar across different cancer types (Extended Data Fig. 1b). We also found that 408 out of the 1211 (33.7%) oncRNAs significantly associated with survival, suggesting that these molecules may have value in predicting patient outcomes (Extended Data Fig. 1c). However, only 30 (2.5%) oncRNAs correlated in expression with early (stages I and II) or late (stages III and IV) stages (Extended Data Fig. 1d). In summary, we discover more than a thousand new cancer-unique RNAs, many of which are not easily traced to the reference human genome.

### Discovery of the first cancer-unique bona fide microRNA

We found some of our novel oncRNAs to be of particular interest. The oncRNA-1 molecule was detected in lung adenocarcinoma samples from both the CPTAC (97/228, 42.5%) and TCGA (324/521, 62.2%) cohorts but was virtually absent in adjacent normal and other healthy tissues (Fig. 3a, Extended Data Fig. 2a). We identified its genomic origin in chr12:116,384,933-116,384,954, an intron of the long non-coding RNA gene *RB11-148B3*.*2* (Fig.3b). Convincingly, we found a very high correlation between the expression of oncRNA-1 and the *RB11-148B*.*2* gene in both TCGA (spearman ρ = 0.81) and CPTAC-3 (spearman ρ = 0.69), supporting that the small RNA is likely processed from this long host transcript (Fig. 3c, Extended Data Fig.2b). At the same time, genes in the chr12q23 region were generally higher expressed in samples with high oncRNA-1 compared to the samples with low oncRNA-1 (Fig. 3d), pointing towards a local de-regulation in that region.

**Figure 3:**
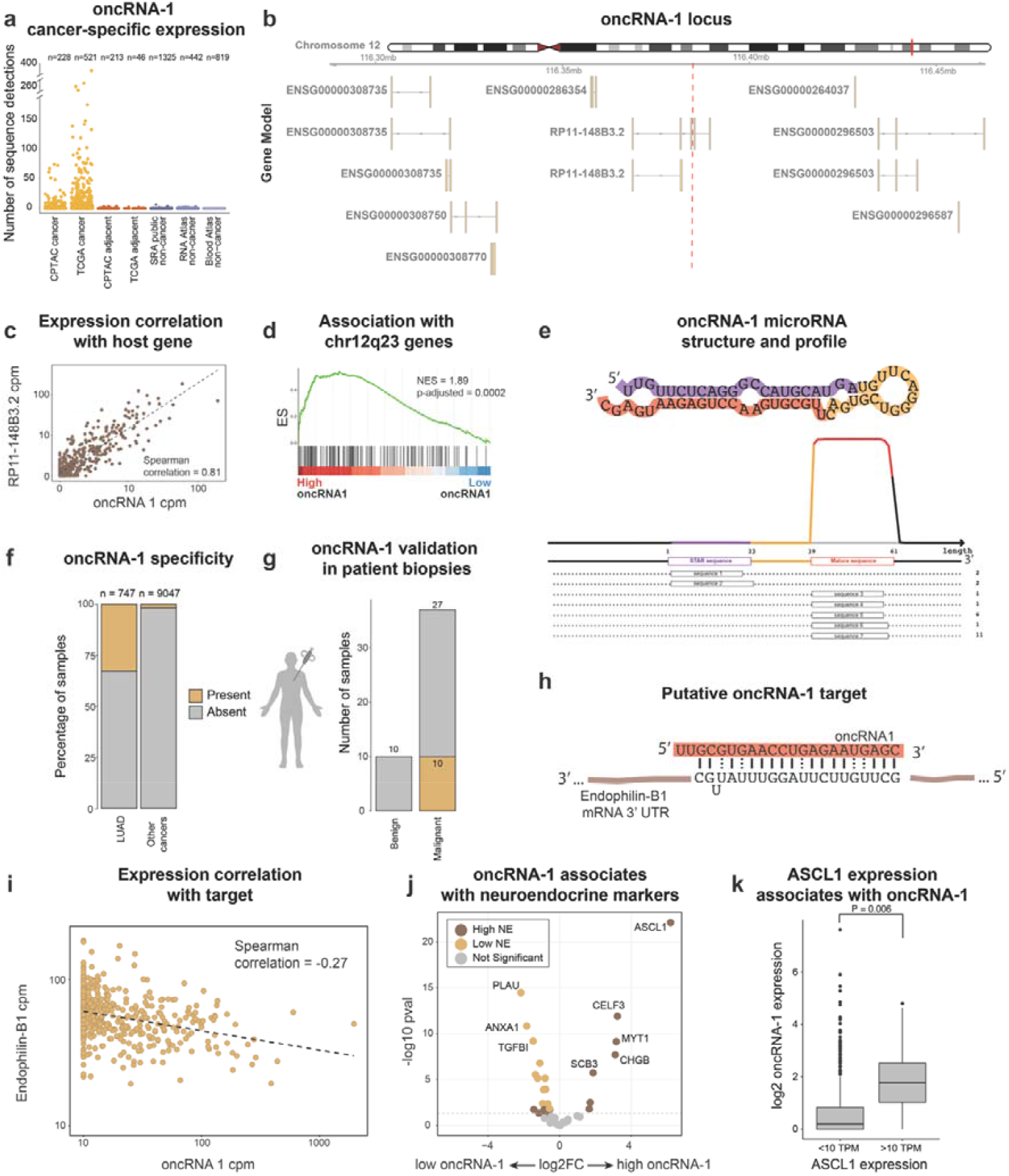
oncRNA-1 is cancer-unique microRNA associated with neuroendocrine differentiation. **a**, oncRNA-1 has abundant expression in lung Adenocarcinoma tumor samples, but is not detected in the adjacent normal tissue and the non-cancer public cohort. **b**, oncRNA-1 is located in the intron of the long non-coding RNARP11-148B3.2, at chr12q23. **c**, oncRNA-1 has high correlation of expression with the host transcript (TCGA; spearman, ρ = 0.81, P = 4.3·10^-121^). **d**, Samples expressing oncRNA-1 show enrichment in genes located at chr12q23 (GSEA analysis, p-adj = 0.0002, NES = 1.89). **e**, oncRNA-1 has many characteristics of a bona fide microRNA, including the secondary structure and the positions of the putative guide (red) and passenger (purple) strand RNAs. **f**, oncRNA-1 is specific to lung adenocarcinoma **g**, The presence of oncRNA-1 was validated in a separate in-house cohort using RT-qPCR. **H**, Endophilin-B1 is a potential target of oncRNA-1. **i**, Expression of oncRNA-1 is negatively correlated with expression of the Endophilin-B1 gene (Spearman’s ρ = −10). 0.27, P = 3.2· ^-10^ **j**, Samples with high oncRNA-1 have high expression of Neuroendocrine markers (differential expression analysis). **k**, Samples with high expression of ASCL1 also express oncRNA-1 (t-test, P = 0.006).

Further analysis revealed that this molecule is transcribed from a MET-element, a mammalian-specific DNA transposon. This particular transposon instance is highly diverged from the consensus sequence (by 17.6% nucleotide substitutions), meaning that it is likely not active. According to miRDeep2^23^ analyses the region around oncRNA-1 adopts an RNA hairpin structure characteristic of microRNAs, and its cleavage pattern is consistent with the canonical microRNA biogenesis pathway (Fig. 3e). Previous studies have shown that transposons with divergent sequences frequently serve as sources for novel microRNAs and other small RNAs^23,24^. This oncRNA appears to be specific to lung adenocarcinoma and is virtually absent in other cancer types. We detect it in only 165 of the 9047 (1.8%) non-lung adenocarcinoma samples in TCGA and CPTAC and 50 out of 2497 (2%) in the control non-cancer atlas (Fig. 3f). Therefore, we conclude that oncRNA-1 is the first cancer-unique miRNA and is only found in lung adenocarcinoma.

### Validation of oncRNA-1 in an independent cohort

Since all our analysis up to this point was conducted on public sequence data from TCGA and CPTAC, we wanted to rule out the possibility of these sequences are computational artifacts of the analysis of these specific cohorts. To do so, we applied TaqMan RT-qPCR to target oncRNA-1 in an in-house lung adenocarcinoma cohort obtained through transthoracic core needle biopsies guided by computed tomography at disease diagnosis. The molecule was present in 10 out of 37 malignant samples and in 0 out of 10 benign (Supplementary Table 3), confirming both its existence and its cancer-specificity using orthogonal methods (Fig. 3g).

### oncRNA-1 associates with neuroendocrine differentiation

Next, we explored the potential function of this microRNA by computationally predicting its targets. Given the low abundance of this molecule, we hypothesized that any function might be similar to an siRNA – efficiently cleaving its targets – rather than performing the subtle repression normally conferred by microRNAs. To test this, we searched for transcripts with extensive complementarity in their 3’ UTRs to the candidate microRNA and identified seven potential targets. Among these, the expression of one target, *Endophilin-B1*, showed a moderate but significant negative correlation to the small RNA in both cohorts (Spearman’s ρ = −0.24, p-adj = 0.007) (Fig. 3h, Fig. 3i). *Endophilin-B1* is known to play a role in autophagy and apoptosis and has been described as a critical tumor suppressor in several cancers^25,26^. These results indicate a potential implication of this cancer-unique microRNA in tumorigenesis.

Additionally, we investigated if oncRNA-1 is associated with any cancer subtype. We performed differential expression analysis on the TCGA and CPTAC datasets (Supplementary Table 4). We compared the 63 samples with zero expression of oncRNA-1 to the 63 samples with highest expression of oncRNA-1, after adjusting for tumor purity (Methods). A total of 5282 protein coding genes and lncRNAs were found to be differentially expressed between the conditions. As expected, *RP11-148B3*.*2*, being the host gene of candidate 1, was the gene with the highest fold-change in TCGA (log2 fold-change = 11.7) (Extended Data Figs. 2c-d). Interestingly, in the samples with high oncRNA-1 expression we observed high fold-changes in many genes usually expressed in neurons. In addition, gene set enrichment analysis revealed association with neuroendocrine cells (Extended Data Fig. 2e). To further test the association with neuroendocrine phenotypes that could be related to cancer^27,28^, we used a set of previously published markers of neuroendocrine differentiation, this set contains 25 genes that are associated with high neuroendocrine differentiation and 25 genes associated with low neuroendocrine differentiation^29^. Strikingly, the results showed a clear trend with most high neuroendocrine markers being up-regulated, especially the *ASCL1* gene, which is a master regulator towards neuroendocrine differentiation. Similarly, most of the negative neuroendocrine markers were down-regulated in the samples with high candidate 1 (Fig. 3j). This association with neuroendocrine differentiation could be partially explained by the fact that *ASCL1* is located in the same arm of chromosome 12, relatively close to the oncRNA-1 and its host gene and are often co-amplified during genomic perturbations in the cancer tissues (Fisher’s Exact Test, P = 0.017, Extended Data Fig. 2f). Finally, we suggest that oncRNA-1 could be used as a predictor of neuroendocrine differentiation in lung adenocarcinoma, since *ASCL1* upregulation is always associated with oncRNA-1 expression (t-test, P = 0.006) (Fig. 3k).

### oncRNA-2 and oncRNA-3 are predictive of patient survival

While oncRNAs are by definition promising biomarkers for detection of cancer, we additionally identified 434 sequences that are not only cancer-unique but are also predictive of patient survival, highlighting their prognostic potential. One such sequence is oncRNA-2, which is present in both lung adenocarcinoma and endometrial carcinoma (Fig. 4a). The presence or absence of oncRNA-2 significantly predicts survival outcomes in patients with endometrial carcinoma across both the TCGA and CPTAC cohorts. Similarly, in lung adenocarcinoma, oncRNA-2 predicts survival in the TCGA cohort, although it does not reach statistical significance in the CPTAC cohort. In all cases, the presence of oncRNA-2 is associated with worse disease outcomes (Fig. 4b). This 25-nucleotide long sequence partially matches an Alu transposon element and maps perfectly to ten genomic locations, many of which fall within genes. We looked at the correlations of all putative host transcripts to the small RNA and found *RHPN2* to be the only gene that consistently correlated strongly (Fig. 4c, Fig. 4d). Notably, this location on chromosome 19 is in an intron of the *RHPN2* gene that transcribes a putative H/ACA Alu snoRNA (Fig. 4c). Shorter RNAs, such as microRNAs, have been shown to be cleaved from snoRNAs^30^.

**Figure 4:**
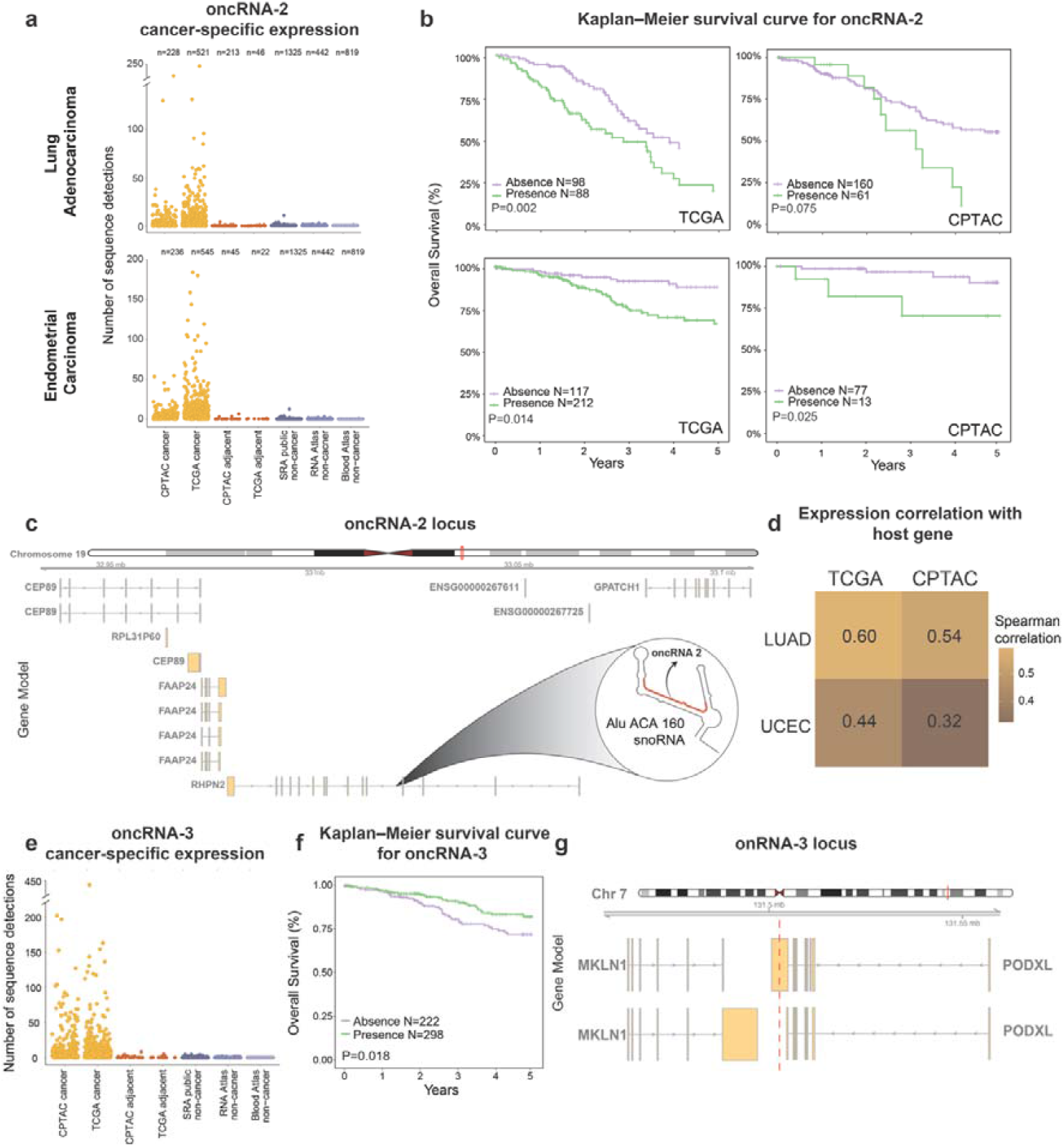
oncRNAs with prognostic value. **a**, oncRNA-2 has abundant expression in lung adenocarcinoma and endometrial carcinoma tumor samples, but is not detected in the adjacent normal tissue and the non-cancer public cohort. **b**, TCGA; Kaplan–Meier plots demonstrate the clinical potential of oncRNA-2 in 186 individuals with LUAD and 329 individuals with UCEC, comparing absence and presence; +, censored observations. Statistical significance was assessed via the log-rank test, with p-values of 0.002 for LUAD and 0.014 for UCEC. CPTAC; Kaplan-Meier plots demonstrate the clinical impact of oncRNA-2 in 221 individuals with LUAD and 90 individuals with UCEC, comparing absence and presence; +, censored observations. Statistical significance was assessed via the log-rank test, with p-values of 0.075 for LUAD and 0.025 for UCEC. **c**, oncRNA-2 originates from an intron of the *RHPN2* gene that gets cleaved into the Alu ACA 160 snoRNA. **d**, Expression of oncRNA-2 correlates well with the host transcript in all cohorts and cancer types. **e**, oncRNA-3 has abundant expression in endometrial carcinoma tumor samples, but is not detected in the adjacent normal tissue and the non-cancer public cohort. **f**, TCGA and CPTAC; Kaplan-Meier plots demonstrate the clinical impact of oncRNA-3 in 520 individuals with UCEC, comparing absence and presence; +, censored observations. Statistical significance was assessed via the log-rank test, with P value of 0.018. **g**, oncRNA-3 originates from the 3’UTR of the *PODXL* gene.

Another example of a cancer-specific molecule associated with patient survival is oncRNA-3. We found this small-RNA expressed in uterine cancer but absent in adjacent healthy tissues from the same patients and in small RNA atlases of healthy tissues (Fig. 4e). Interestingly, oncRNA-3 appears to be associated with better survival outcomes (Fig. 4f). oncRNA-3 maps to an exon of the mRNA encoding *PODXL* (Fig. 4g), a protein previously implicated in cancer progression. It is well-established that structures in mRNAs can be cleaved into small RNAs, effectively leading to gene silencing^22^. Given that full-length *podxl* mRNA has been linked to poor survival in specific cancer types^31–33^, this suggests a mechanism where cleavage of the transcript may generate the small RNA, silencing the gene and attenuating its aberrant overexpression. Profiling both the full-length *podxl* mRNA and the derived cancer-specific small RNA could thus provide valuable insights into patient prognosis.

### oncRNA-4 originates from an ancestral DNA sequence that has been lost in most humans

One of the advantages of our unbiased reference-free method is that we can discover RNAs that are not present in the human reference genome. An example of such a case is oncRNA-4, a cancer-unique molecule that was found in endometrial carcinoma (Fig. 5a). We examined the expression of the molecule in all cancer types that are available in TCGA and discovered that it has high expression in two other female cancers, breast cancer and ovarian cancer (Fig. 5b). We found that oncRNA-4 is part of a 46 nucleotides long genomic variant, an evolutionarily ancestral DNA sequence that is present in most studied primates but is absent in the majority of human individuals (Fig. 5c). More specifically, the variant was only present in around ~25% of the samples from the Human Pangenome Atlas^14^ (Fig. 5d). In the endometrial carcinoma samples, the genomic variant was present in 96% of the patients that expressed oncRNA-4, but only 14% of the patients that did not express the oncRNA, according to the matched DNA sequencing data. We also found the genomic variant in the genomic sequences from blood samples, confirming that it is a not a de novo cancer mutation (Fig. 5e). Around 40% of samples examined showed both the variant and the alternative allele suggesting that these patients were heterozygous, whereas 10% exhibited only the variant (Fig. 5e). At the same time, the expression of the small RNA correlated with the presence of the variant in the DNA and was also dependent on whether the patient is homozygous or heterozygous (Fig. 5f). In summary, we report a small RNA that is specific to female cancers and originates from an ancestral part of the human genome that has been lost in most human individuals.

**Figure 5:**
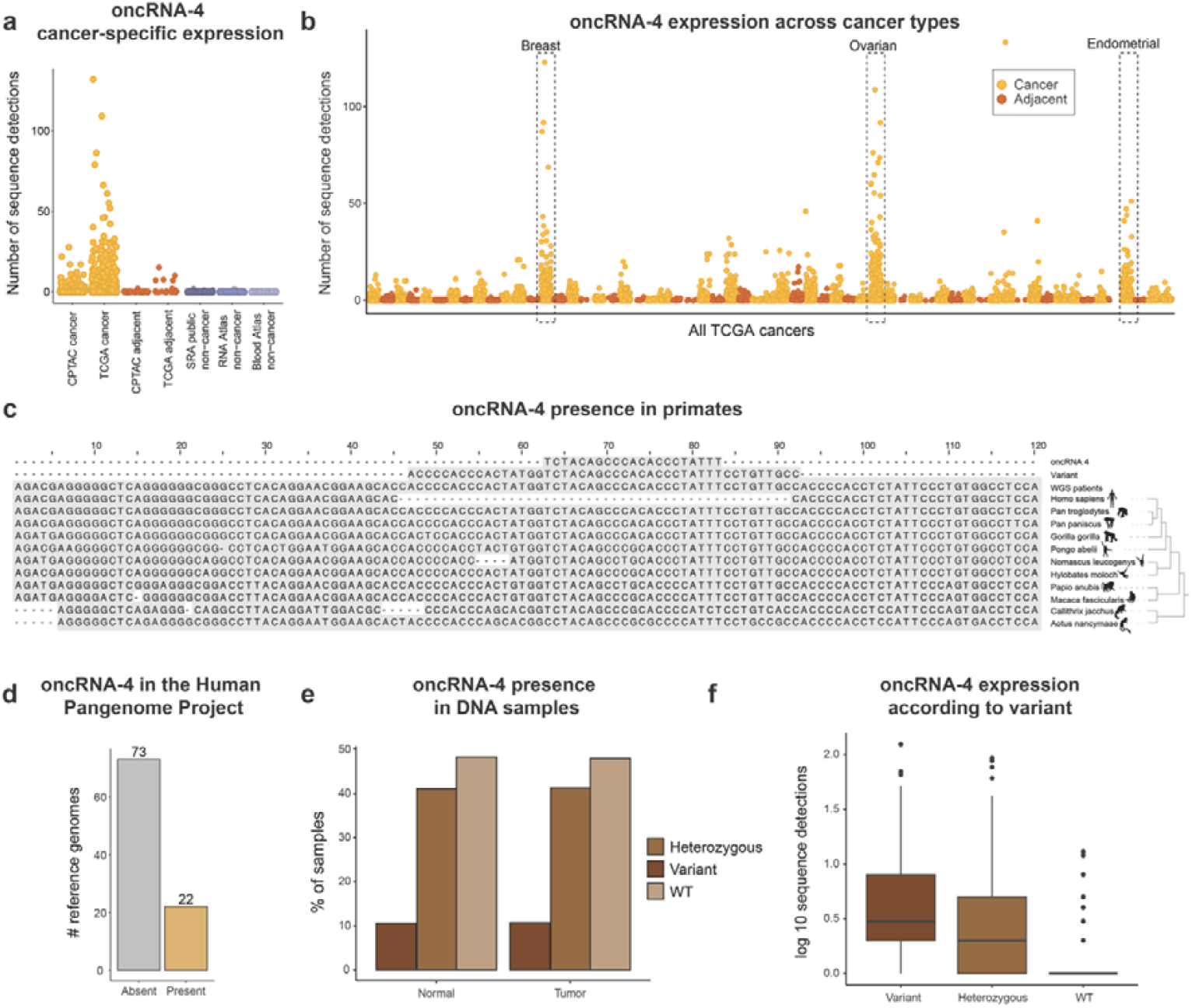
oncRNA-4 does not map to the human reference genome. **a**, oncRNA-4 is expressed in endometrial carcinoma tumor samples, but is not detected in the adjacent normal tissue and the non-cancer public cohort. **b**, oncRNA-4 is most highly expressed in three female cancers: breast cancer, ovarian cancer and endometrial cancer. **c**, The genomic variant that oncRNA-4 is expressed from is absent from the human reference genome but conserved in the reference genomes of most primates. **d**, The variant is present in ~25% of samples from the human pangenome project. **e**, The variant that includes oncRNA-4 is found in blood and tumor DNA in similar proportions. **f**, The expression of oncRNA-4 correlates with presence of the variant in DNA.

## Discussion

In this study, we developed and applied an unbiased, reference-free computational approach to identify cancer-unique oncRNAs across multiple cancer types. Since the method does not require genome mapping, it can discover sequences that are not part of existing annotations or that are not even present in the human reference genome. Although here we only applied our method to analyze cancer data, we expect that the method is also applicable to any situation where two conditions are compared and the use of annotations or reference genomes is not possible. For example, other diseases (especially the ones that are associated with higher genomic instability), poorly annotated model organisms or even complex environmental metatranscriptomic samples^35,36^.

Previous efforts to identify cancer-associated molecules have largely centered on identifying RNAs that are differentially expressed in the diseased condition^37,38^. These studies commonly employ targeted approaches aimed at known small RNAs. While such methods can be informative, their reliability is often constrained by assay reproducibility across experimental conditions. Moreover, these approaches typically rely on distributional comparisons, which may fail to capture outlier cases, thereby limiting their sensitivity in heterogeneous patient populations. In contrast, oncRNAs are binary biomarkers that can classify samples simply by their presence or absence. They therefore have the potential to be highly useful biomarkers.

We successfully discovered more than a thousand oncRNAs across five cancer types and focused on four molecules of interest. At the same time, we delve into the biogenesis oncRNAs and show examples of what these molecules are. Notably, one of them, oncRNA-1 originates from the intron of a long non-coding RNA, a region that folds in a microRNA hairpin structure. This, to our knowledge, is the first case of cancer-unique bona fide microRNA. Other oncRNAs originate from the 3’UTRs of mRNAs, for example oncRNA-3. Additionally, we show that a substantial number of oncRNAs is associated with patient survival and can thus have prognostic potential, for example oncRNA-2 and oncRNA-3. In addition, we discover oncRNAs that do not map to the human reference genome. As shown by oncRNA-4, these molecules can be of particular interest and can be used as keyholes into cancer biology.

Our results indicate that oncRNAs tend to be highly specific to individual cancer types, with limited overlap observed between different cancers. For instance, we found only modest overlap between the two non-small cell lung cancers, LUAD and LUSC, which share a similar tissue of origin. This specificity implies that the mechanisms driving the production of oncRNAs are distinct and potentially reflective of the unique genomic and epigenomic landscapes of each cancer type. This observation is consistent with previous studies that emphasize the heterogeneity of cancer, both within and between tumor types.

One important takeaway message from our study is that only a small percentage of the oncRNAs found in the TCGA cohort overlapped with the oncRNAs found in the CPTAC cohort. This could be due to intra-cohort variability but also highlights the need for careful validation of oncRNA biomarkers. Although the use of two cohorts is not always possible, with the increasing amount of publicly available data it is becoming more and more of a viable strategy. We recognize that our approach is very strict and we expect to have missed many bona fide oncRNAs. For instance, cases of sample swaps or human tissues with undiagnosed cancer in our supposed non-cancer controls could make oncRNAs appear to not be unique to cancer, and therefore missed by our approach. Therefore, we are likely just discovering the tip of the iceberg in our current study.

A recent study presented in a preprint reported ~260 000 novel oncRNAs across 32 cancer types, some of which were found to be functionally active in vivo or in cellular systems^39^. The oncRNA molecules that we present here have passed more stringent filters, given that they were discovered to be cancer-unique in two independent cohorts, TCGA and CPTAC. It will be interesting to see what the overlap is between the two sets of sequences, and the future emergence of more oncRNAs might necessitate the development of a database where the sequences can be categorized and stratified according to their confidence levels and their functional and clinical relevance.

In conclusion, we here deliver a new method for unbiased reference-free identification of small RNAs, the hundreds of oncRNA molecules that we identified with it as a resource for the community and biological insights into the biogenesis and role of these molecules in cancer.

## Methods

### Data acquisition

Small-RNA sequencing data were downloaded from Genomic Data Commons for both the TCGA and CPTAC cohorts. Only samples marked as “Primary Tumor” or “Adjacent Tissue Normal” were considered.

| Cancer type | TCGA Project ID | CPTAC-3 Project Name |
| --- | --- | --- |
| Lung Adenocarcinoma | TCGA-LUAD | Lung; Adenomas and Adenocarcinomas |
| Lung squamous cell carcinoma | TCGA-LUSC | Lung; Squamous Cell Neoplasms |
| Pancreatic Adenocarcinoma | TCGA-PAAD | Pancreas |
| Kidney Adenocarcinoma | TCGA-KIRC, TCGA-KIRP, TCGA-KICH | Kidney |
| Endometrial carcinoma | TCGA-UCEC | Uterus |

### The pipeline

Raw sequencing data were downloaded as BAM files using the gdc_client tool version 2.3 and aggregated by condition samtools merge version 1.20^40^. These BAM files were converted to FASTQ format using samtools fastq. Quality control of the resulting FASTQ files was performed with mirtrace qc version 1.0.1^41^, using the option to retain uncollapsed FASTA outputs. K-mer counting was then carried out on the uncollapsed FASTA files using KMC version 3.2.4^42^.

This preprocessing pipeline was executed separately for tumor and adjacent (non-tumor) tissue samples. Subsequent analysis was conducted in R version 4.4.1. k-mers enriched in cancer samples were identified based on the following criteria:

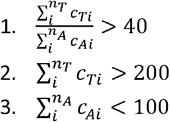

Where c_T_ and c_A_ the counts for each tumor or adjacent sample respectively and n_T,_ n_A_ the number of tumor and adjacent samples.

Only k-mers satisfying these thresholds in both TCGA and CPTAC cohorts were retained. Their expression levels were then quantified in an independent cohort of non-cancer samples. K-mers with more than 200 counts in this control cohort were filtered out. Finally, overlapping and offset k-mers were merged using dekupl-mergeTags^43^ to generate a curated list of candidate cancer-specific sequences. The final sequences were also tested against known adapter and primer sequences to exclude artifacts with *bash grep*.

### oncRNA annotation and mapping

oncRNAs were mapped to the human genome (GENCODE release 48) with bowtie version 1.3.1^44^ with the -v 0 and -v 1 --all and --strata options. Genome annotations were added with a custom script using the bedtools version 2.31^45^ intersect function.

### microRNA prediction

microRNA hairpin predictions were performed with miRDeep2 version 0.1.3^23^.

### Target predictions

Small RNA target prediction was conducted with RIsearch 2.1^46^ on the 3’UTRs of ENSEMBL transcripts with a minimum seed length of 15nt.

### qPCR validations of oncRNA-1

Total RNA, including small RNAs, was extracted from transthoracic lung biopsies and lung adenocarcinoma surgical specimens using the ReliaPrep miRNA Cell and Tissue Miniprep System (Promega).

The oncRNA-1 sequence (TTGCGTGAACCTGAGAATGAGC) was submitted for custom TaqMan Small RNA Assay (ThermoFisher Scientific) and validated using TaqMan RT-qPCR with U6 snRNA (#001973, ThermoFisher Scientific) as an internal control. A total of 10 ng of RNA was used as input for each reaction. Detection was called using a cycle threshold (Ct) cutoff < 35. Ct-values for U6 were stable around 20 across all samples.

### Sample acquisition of in-house cohort

A total of 47 samples were collected, corresponding to 10 patients with benign disease and 37 patients with lung adenocarcinoma. All benign samples (including hamartoma, inflammation and sarcoidosis), were acquired through transthoracic core needle biopsies. Of the malignant samples, all confirmed as lung adenocarcinoma, two were from surgically resected material and forty from transthoracic core needle biopsies. Samples were directly put in CELLBANKER 2 solution followed by storage in −80C until RNA extraction. All samples were approved for analysis by the regional ethical review board in Stockholm (DNR 2018/1246-32 and DNR 2018/1246-32/1). Written informed consent was obtained from all patients analyzed in this study.

### Bioinformatic analyses

TCGA transcriptomic data was downloaded with TCGAbiolinks version 2.32.0^47^. All differential expression analyses were carried out with DESeq2 version 1.44^48^. Gene set enrichment analysis was performed with fgsea version 1.30^49^. Survival regression analysis was performed using the R package “survival” version 3.8-3. Stratification of patients was made as follows: Absence: 0 counts, Presence: > 5 counts.

### Copy number amplification analysis

Allele-specific copy number segments obtained from ASCAT3 (TCGAbiolinks, version 2.34.0) were analyzed for samples in the top and bottom candidate expression groups (N = 60 and N = 62, respectively). Each sample was classified according to the copy number status of the segment fully encompassing the host gene ENSG00000258346. Segments were categorized as follows: total copies < 2, Loss; total copies > 2, Gain; total copies > 4, Amplification; and total copies > 10, High-level amplification. Loss of heterozygosity (LOH) was defined as cases where the number of major alleles equaled the total copy number. Statistical significance between top and bottom groups was assessed using Fisher’s exact test for contingency tables.

### Tumor purity adjustments

Expression of oncRNA-1 was adjusted based on tumor purity. The tumor purity estimates where retrieved by using “ESTIMATE” from TCGAtumor_purity() function of TCGAbiolinks version 2.32.0.

### Multiple sequence alignment

Multiple sequence alignment of the primate sequences was performed with Clustal Omega online tool^50^.

## Acknowledgements

M.R.F and P.K. acknowledge funding from the VR Consolidator Grant no. 2022-03953 ‘InSync’ and the Cancerfonden project grant 243710Pj ‘Discovery and characterization of cancer-specific small RNAs’. The results shown here are in part based upon data generated by the TCGA Research Network: https://www.cancer.gov/tcga. Data used in this publication were generated by the National Cancer Institute Clinical Proteomic Tumor Analysis Consortium (CPTAC). The computations and data storage were enabled by resources provided by the National Academic Infrastructure for Supercomputing in Sweden (NAISS), partially funded by the Swedish Research Council through grant agreement no. 2022-06725.

## Author contributions

M.R.F, B.F and P.K conceived, designed and planned the study. P.K performed all of computational analyses, except as specified here. K.D performed copy-number variant analyses. M.A.U performed the initial oncRNA annotation analyses. S.E, P.H and A.T designed and performed the qPCR validation studies based on in-house cohorts. M.R.F supervised computational analyses. P.K and M.R.F wrote the manuscript with input from all authors.

